# Alternate isoforms of IRF7 Differentially Regulate Interferon Expression to Tune Response to Viral Infection

**DOI:** 10.1101/2025.03.10.642367

**Authors:** Asmita Panthi, Max B. Ferretti, Olivia Howard, Swechha Mainali Pokharel, Rhiannon McCracken, Mathieu Quesnel-Vallieres, Qin Li, Sara Cherry, Kristen W. Lynch

**Affiliations:** Department of Biochemistry and Biophysics, University of Pennsylvania, Philadelphia, PA, 19104, USA; Pharmacology Graduate Group, University of Pennsylvania, Philadelphia, PA, 19104, USA; Department of Pathology and Laboratory Medicine, University of Pennsylvania, Philadelphia, PA, 19104, USA; Biochemistry, Biophysics and Chemical Biology Graduate Group, University of Pennsylvania, Philadelphia, PA, 19104, USA; Department of Genetics, Perelman School of Medicine, University of Pennsylvania, Philadelphia, PA, 19104, USA

**Author notes:** Correspondence should be addressed to K.W.L.

**Keywords:** Alternative Splicing, IRF7, Interferon, Transcription Activation, Innate Immunity

## Abstract

Interferon Regulatory Factor 7 (IRF7), and its homologue IRF3, are master transcriptional regulators of the innate immune response. IRF7 binds to promoters of interferon β (IFNβ) and several IFNαs as a homodimer or as a heterodimer with IRF3 to drive expression of these type I IFNs, which in turn activate downstream signaling pathways to promote expression of antiviral genes. Here we demonstrate that alternative splicing of the first intron within the coding region of IRF7 is highly regulated across immune tissues and in response to immunologic signals including viral infection. Retention of this intron generates an alternative translation start site, resulting in a N-terminally extended form of the protein (exIRF7) with distinct function from the canonical version of IRF7 (cIRF7). We find that exIRF7 uniquely activates a gene expression program, including IFNβ, in response to innate immune triggers. Mechanistically, this enhanced activity of exIRF7 relative to cIRF7 is through increased homodimerization and association with IRF3 on DNA. Furthermore, the enhanced transcriptional activity of exIRF7 controls viral infection to a greater extent than cIRF7, demonstrating that alternative splicing of IRF7 is a previously unrecognized mechanism used by cells to tune the interferon response to control viral infections and other immune challenges.

**Highlights:**

- Intron retention in the human IRF7 gene generates a distinct protein isoform that differs in the N-terminus.
- IRF7 intron retention is regulated in a stimuli- and cell-type specific manner.
- The extended version of IRF7, produced by intron retention, exhibits enhanced transcriptional activation of type I interferon genes.
- Cells expressing the extended version of IRF7 are more resistant to viral infection.

## Introduction

Alternative splicing is a mechanism of gene regulation that broadly controls protein identity through the differential inclusion or skipping of segments of pre-mRNA, which can alter the encoded protein. Such alternative splicing is often regulated in response to cellular signals to enable cells to modulate protein function in response to extracellular cues^1–4^. In particular, work in recent years has revealed widespread changes in alternative splicing in response to viral infection and innate immune signaling^5–9^. Despite this recent progress, however, we have a poor understanding of the extent to which alternative splicing plays a role in tuning the interferon response, which functions as a central component of innate immunity.

Interferon Regulatory Factor 7 (IRF7), and its homologue IRF3, are master transcriptional regulators of the innate immune response that drive expression of type I interferon genes (IFN-I) including IFNβ and IFNαs ^10–12^. Type I interferons (IFN-I), in turn, are secreted and bind their receptor in an autocrine and paracrine fashion to activate JAK-STAT signaling, thereby inducing the expression of numerous Interferon Stimulated genes (ISGs), many of which inhibit viral replication and activate cellular pathways to combat infection^13,14^. IFNs and ISGs also play a central role in tuning inflammation and susceptibility to autoimmune diseases^12^.

IRF7 and IRF3 are both comprised of an N-terminal DNA binding domain (DBD), a central transactivation domain (TAD), and a C-terminal region that undergoes extensive post-translational modifications to regulate dimerization, localization and function^12^. In response to viral infection, or detection of pathogen-associated molecular patterns (PAMPs) by pattern recognition receptors (PRRs), IRF7 and IRF3 are activated by phosphorylation, which induces dimerization and nuclear translocation^10,12^. Once in the nucleus, IRF7 binds to a range of related but distinct DNA sequences, including interferon regulatory factor-binding elements (IRFEs) upstream of IFN-I mRNAs and interferon-stimulated response elements (ISREs) upstream of ISG mRNAs^15^. Through these elements, IRF7 activates transcription of type I IFN mRNAs in response to viral PAMPS, and also promotes expression of ISG mRNAs both directly and via IFN-induced JAK-STAT signaling^15,16^. Of note, IRF7 and IRF3 are typically thought to function as a heterodimer with one another, but can also function as homodimers. IRF7 and IRF3 also associate with additional transcription factors, most notably NF-kB and ATF-2/cJun to activate the IFNβ promoter^10,15–18^.

IRF3 is ubiquitously expressed across tissues and is regulated solely by phosphorylation^19^. By contrast, IRF7 expression is more variable: in most cell types IRF7 is expressed at only low levels under basal conditions, but it is constitutively expressed in barrier epithelial cells and some lymphoid populations to promote initial induction of IFN-I^20,21^. Regardless of basal expression of IRF7, in all cell types IRF7 is strongly induced by IFN-I signaling to form a positive feedback loop that amplifies IFN-I pathway activation^12,15,22^. In mouse models, loss of IRF7 has a greater impact on IFN-I expression and anti-viral immunity than loss of IRF3^11,23^. Moreover, humans with loss of function mutations in IRF7 are highly susceptible to viral infections^11,24,25^ and autoimmune diseases^12,26^. Therefore, control of IRF7 expression and activity is essential for tuning proper immune function and productive anti-viral defense.

We have previously demonstrated that both adaptive immune signaling and viral infection induce changes in splicing that promote productive immune responses^6,27–29^. Therefore, we explored genes involved in innate immunity for regulated alternatively splice forms that may impact innate signaling. Strikingly, we find that inclusion of the first intron of the IRF7 coding region is highly variable across immune cell types and is tightly regulated in response to a range of immune stimuli. Inclusion of this intron alters the translation start site and results in a novel 19 amino acid N-terminal extension adjacent to the DNA binding domain of this key transcription factor. We find that the extended IRF7 (exIRF7) isoform exhibits enhanced activity on the promoters of type I IFNs, particularly IFNB1 which encodes IFNβ. This enhanced activity is due to an increased homodimerization and heterodimerization with IRF3 on DNA and results in increased anti-viral immunity through increased IFN-I production. Together, these data reveal a previously unrecognized mechanism by which the interferon response is regulated through alternative splicing of IRF7.

## Results

### Alternative splicing of human IRF7 is regulated in a tissue- and stimulation-dependent manner

Given the precedent that adaptive immune signaling regulates alternative splicing of genes which tune the immune response^1,3,6,27,28,30^, we explored alternative splicing of canonical innate immune factors. We focused on IRF7 as this transcription factor plays a central role in the interferon response and in mouse macrophages TNF-α treatment causes downregulation of IRF7 protein via retention of *IRF7* intron 4, which generates a premature stop codon^31^ (Fig S1A). The initial cloning and characterization of the *IRF7* gene in humans also reported an isoform in which the first intron after the translation start site is retained^32,33^. Retention of this first intron is predicted to result in translation from an in-frame intronic start site to yield a unique extended N-terminus (Fig 1A, exIRF7, also called IRF7D and IRF7H in previous literature^32,33^); however the protein product of this extended isoform was not characterized in detail as data at the time suggested this extended isoform was not highly expressed in cells tested and did not vary upon viral infection^32,33^.

**Figure 1.**
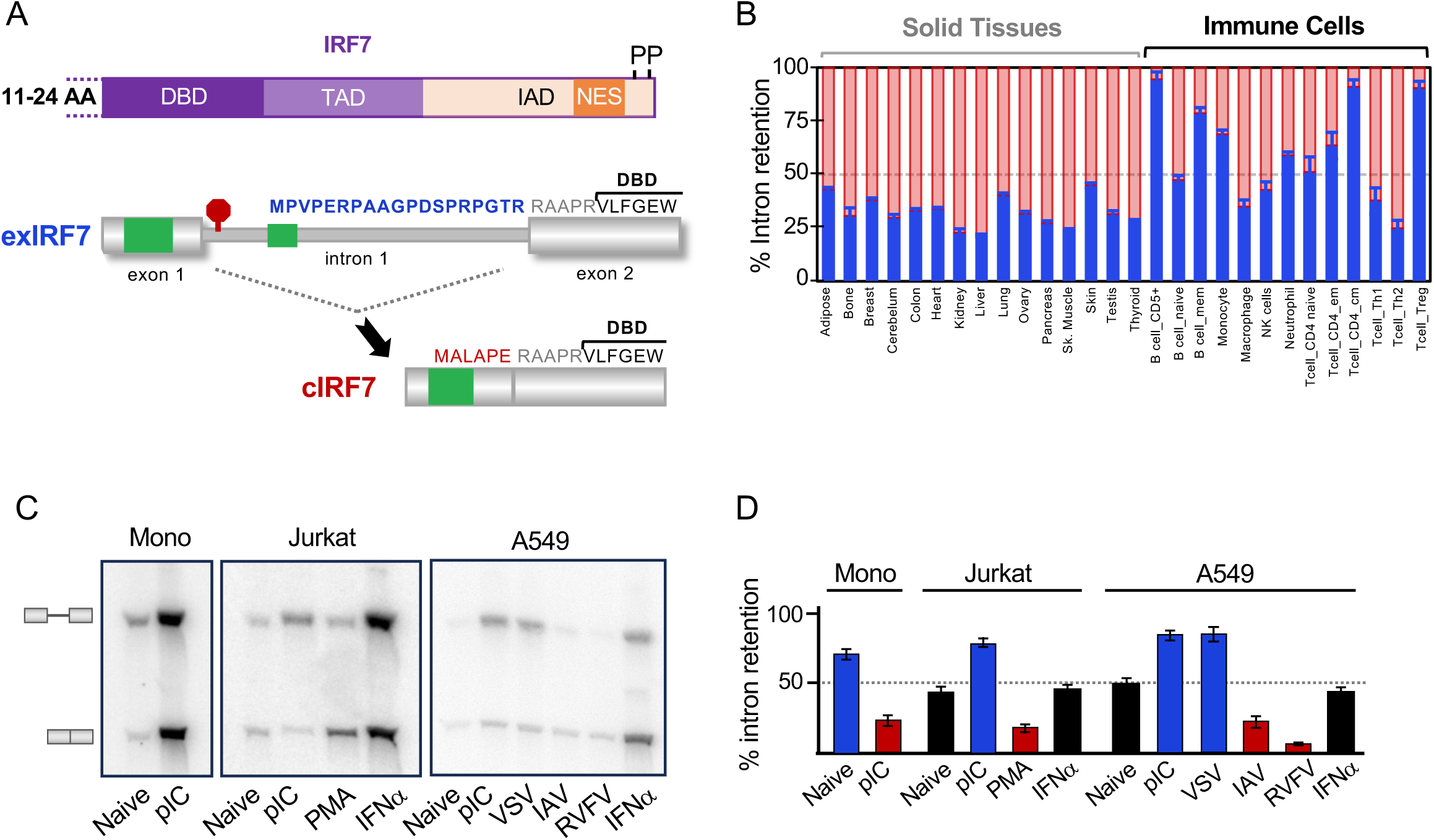
Alternative splicing of human IRF7 Intron 1 is regulated in a tissue-specific and stimulation-dependent manner. (A) Schematic of domains of the IRF7 protein (top): DNA binding domain (DBD), trans-activation domain (TAD), IRF-association domain (IAD), and nuclear export signal (NES). (bottom) Schematics of the alternatively spliced variants indicating the translation start sites in exon 1 and retained intron (green boxes), and stop codon in intron 1 (red octogon). Isoform-specific amino acids preceding the DBD are indicated in blue for exIRF7 and red for cIRF7. (B) Plot of IRF7 intron 1 retention across human cell types from public GTEx data. Blue bars indicate percent of total IRF7 expression that retains intron 1. Red area indicates percent of cIRF7 isoform. (C) RT-PCR analysis of IRF7 intron 1 alternative splicing in various human cell types under different stimulation conditions. (D) Quantification of the RT-PCR data in panel C. Conditions that preferentially express exIRF7 are shown in blue, conditions that preferentially express cIRF7 are in red. Black bars indicate near equal expression of both isoforms.

Strikingly, using publicly available transcriptomic data^34^, we observe marked variability in the retention of the first intron across primary human immune cell types, ranging from almost full inclusion in regulatory and memory T and B cells to predominantly spliced in macrophages and differentiated CD4+ T cells (Th1, Th2) (Fig 1B, blue bars). By contrast, intron 1 exhibits low and consistent inclusion in non-immune cell types (Fig 1B) and intron 4 exhibits low and consistent inclusion across all cell types (Fig S1B). Importantly regulation of intron 1 is independent of transcript abundance or retention of intron 4 (Fig S1B) suggesting a distinct mode of control.

As monocytes differentiate to macrophages during infection and inflammation, the difference in *IRF7* intron 1 retention between monocytes and macrophages suggests that this splicing event is regulated in response to immune triggers. We first confirmed the preferential retention of *IRF7* intron 1 in purified human monocytes by RT-PCR (Fig 1C,D). Consistent with immune-induced regulation of splicing, treatment of cultured human monocytes with poly-inosine:cytosine (pIC), a mimic of dsRNA as sensed during viral infection, leads to a clear decrease in retention of intron 1 (Fig 1C,D). By contrast, treatment of CD4 T cell-like Jurkat cells or A549 lung epithelial cells with pIC leads to a dramatic increase in *IRF7* intron 1 retention, resulting in more exIRF7 mRNA (Fig 1C,D). Interestingly, infection of A549 cells with several RNA viruses has the opposite effect, increasing splicing rather than promoting intron retention (Fig 1C,D), suggesting that these viruses have some activity that counters the dsRNA-induced response. Treatment of Jurkat cells with PMA, which activates Ras signaling in a manner analogous to T cell signaling, also increases splicing of *IRF7* intron 1 (Fig 1C,D).

Given that pIC, PMA and viral infection all induce the transcription of *IRF7* (Fig 1C), but have distinct impacts on splicing (Fig 1C,D), we conclude that changes in transcript abundance does not drive the splicing changes in *IRF7* intron 1. Further confirmation of the lack of correlation between transcription and splicing is evident in the fact that treatment of both Jurkat and A549 with IFNα, which strongly induces expression of IRF7, results in no change in IRF7 splicing (Fig 1D). Together these data demonstrate that the splicing of IRF7 intron 1 is tightly and directly regulated in a signal-dependent manner across many different cell types and immune triggers, independently of IRF7 transcription. This signal-induced regulation of intron retention is specific for IRF7 intron 1, as we observe no change in intron 4 splicing in either Jurkat or A549 cells in response to any stimuli or virus tested (Fig S1C).

### Isoforms generated via alternative splicing of IRF7 intron 1 exhibit distinct transcriptional activities

Since we observed regulated changes in *IRF7* intron 1 inclusion we focused our mechanistic studies on this alternative splice form. Moreover, unlike most examples of intron retention, which inhibit protein production through the addition of premature stop codons, both isoforms generated through alternative splicing of *IRF7* intron 1 yield an open reading frame differing only in the N-terminal 6 or 19 amino acids, immediately adjacent to the DNA binding domain. This led us to hypothesize that there may be altered transcriptional activity different between the two IRF7 isoforms.

As a first step to determining differences in function between cIRF7 and exIRF7 we used CRISPR gene editing to generate Jurkat clones that can only express one isoform of IRF7. We made targeted cuts outside of the introns flanking the regulated first coding intron and used homology repair to replace this region with either a fusion of the flanking exons (thus modeling splicing and expression of cIRF7), or a version of the gene carrying two base changes at the 5’ splice site of the intron which prevents splicing (thus enforcing expression of exIRF7). The 5’ splice site mutation used does not cause any change in the encoded amino acid, or the presence or location of the start and stop codons (Fig 2A, S2A), thus retaining all the translation information and potential regulation of the native gene. We confirmed two clones of each genotype by Sanger sequencing and demonstrated by RT-PCR that these clones express only the predicted RNA isoform (Fig 2B, S2B-C). Importantly, each of these clones express similar levels of total IRF7 protein (Fig 2B) upon stimulation with IFNα, confirming that the exIRF7 RNA isoform is translated into a stable protein. As a further control for subsequent experiments, we also generated IRF7 knockout (KO) cell lines by deleting the third coding exon by CRISPR, generating a premature termination codon (Fig S2B,D), and we confirmed loss of protein expression by Western blot (Fig 2B, bottom).

**Figure 2.**
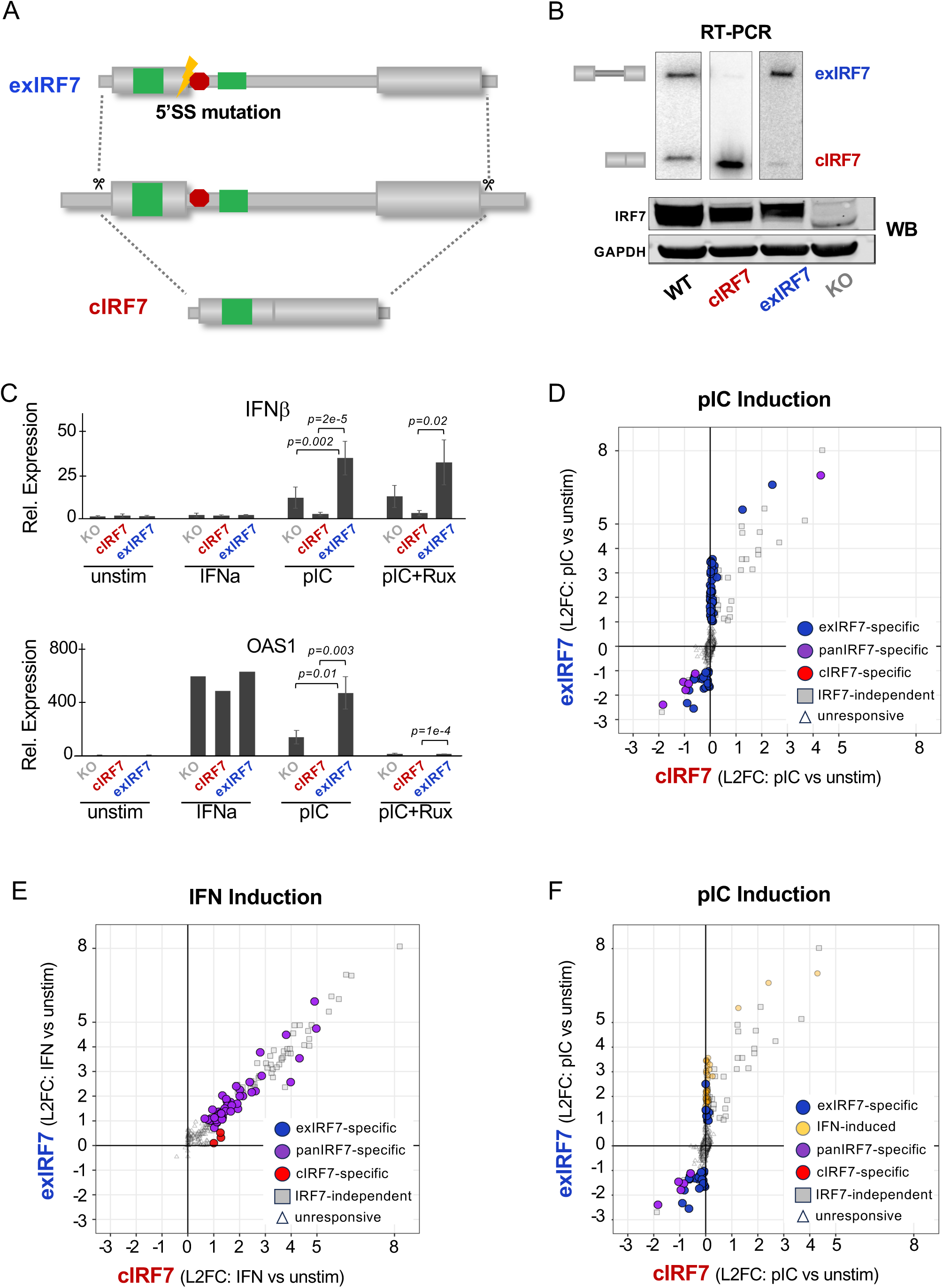
Isoforms generated through alternative splicing of IRF7 intron 1 exhibit distinct transcriptional activities in response to innate immune activation. (**A**) Design of CRISPR-Cas9 genome editing to create cell lines expressing either cIRF7 or exIRF7. (**B**) Validation of CRISPR tools at both RNA and protein levels: (top) RT-PCR confirms the expression of IRF7 isoforms at the RNA level compared to WT cells. (bottom) Western blot analysis reveals the protein expression levels of exIRF7 and cIRF7 relative to WT and IRF7 KO clones. (**C**) RT-qPCR analysis of selected differentially expressed genes (DEGs) from RNA-seq (n=3, mean ± SD) under conditions of IFNα and pIC, with and without ruxolitinib treatment. Differences that are significant are indicated (p<0.05, unpaired student t-test). (**D-F**) Scatter plots based on RNA-seq data showing differentially expressed genes (DEGs) in exIRF7 and cIRF7 compared to IRF7 KO cells after induction with pIC (10 µg/ml for 6 hours, **D, F**) or IFNα (500 U/ml for 6 hours, **E**). Panel F is the same as panel D but with different gene groupings indicated. Significant DEGs were identified with a base read minimum of 50, a minimum log2-fold change of 1 and a p-value threshold of 0.05.

We first tested how each IRF7 isoform impacted expression of type I interferons, as these are the canonical targets of IRF7 downstream of PRR engagement. As expected, we observed modest induction of IFNβ upon stimulation with polyIC in IRF7KO cells which express IRF3. Strikingly, we observe enhanced IFNβ induction in exIRF7 cells, and suppression in cIRF7 cells relative to IRF7-KO cells (Fig 2C). Furthermore, by both qPCR (Fig 2C, S3A) and RNA-Seq (Fig 2D) across two independent clones of each genotype, we identify 73 genes that are significantly upregulated by pIC treatment in the exIRF7 cells but unchanged in the cIRF7 or IRF7-KO cells, 59% of which are known ISGs. A general lack of responsiveness to pIC in cIRF7 expressing cells is also observed by principal component analysis, which reveals that the pIC-treated cIRF7 cells cluster with unstimulated exIRF7 and IRF7-KO cells (Fig S3B).

By contrast to this pIC-induced expression, direct treatment of the CRISPR clones with type I IFN identifies ∼50 ISGs which are similarly induced in cIRF7 and exIRF7-expressing cells relative to the IRF7-KO cells (Fig 2E, panIRF7-specific; Fig S3A, STAT2). This panIRF7 activation is consistent with reports that IRF7 can cooperate or act in parallel with the JAK-STAT induced ISGF3 complex to enhance transcription of some ISGs^15,16^, and importantly confirms that cIRF7 in our cells retains transcriptional activity under some conditions. As expected, we also observe a broad program of ISG induction by IFN-I that is independent of IRF7 (i.e. genes where induction in the IRF7-KO cells is similar to those in the cIRF7/exIRF7 expressing cells), presumably through the sole activity of ISGF3 (Fig 2E, IRF7-independent; Fig 2C, OAS1).

As further evidence that the ISGs induced by pIC in exIRF7 cells are a secondary effect of exIRF7-induced IFNβ, 64% of the genes induced by pIC in the exIRF7-expressing cells overlap with those induced by IFNα in the IRF7-KO cells (Fig 2F, yellow dots). Moreover, treatment of cells with the JAK inhibitor ruxolitinib, blocks JAK-STAT signaling downstream of IFN-I receptor engagement, abolishes the pIC-induced expression of the ISGs OAS1 and STAT2 in the exIRF7 expressing cells (Fig 2C, S3A), but has no impact on the exIRF7-specific pIC-induction of IFNβ, or genes such as NFKBIA, which are not induced by IFNα treatment (Fig 2C, S3A). Together, these data suggest that exIRF7 uniquely regulates a subset of genes upon pIC activation (Fig 2F, blue dots), which critically includes IFNβ, that subsequently activates an ISG response.

Of note, the set of genes uniquely induced in exIRF7 cells by pIC also includes several repressors of transcriptional activation (NFKBIA, CXXC4, DAB2IP; Fig S3A and Table S1) which may explain the down-regulated genes observed in pIC treated cells (Fig 2F, blue dots with negative values). Further evidence that the down-regulated genes are indirectly controlled by IRF7 is that genes that are repressed upon pIC in the exIRF7 cells mostly lack an ISRE consensus (GAAANNGAAA) near the transcription start site (TSS) (Fig S4A-C, light blue). Indeed, the pattern of predicted ISREs in this set mirrors that of non-responsive genes (Fig S4A-C, gray), and in the few repressed genes that do score as having an ISRE, there is typically only a single copy which is a poor match to the consensus (Fig S4D). By contrast, the majority of genes that are enhanced by either isoform of IRF7 in response to IFNα (pan-IRF7 enhanced) or enhanced by exIRF7 in response to pIC, have 2-4 strong matches to the ISRE consensus motifs within 200 nucleotides of the TSS (Fig S4A-D, purple).

Finally, to determine if IRF7 isoform-specific gene expression is also observed in other cell types, we engineered human A549 lung epithelial cells to express single isoforms of IRF7 (Fig S5A-B). As we were unable to use the same CRISPR repair strategy to force IRF7 isoform expression in A549 cells due to inefficiency of homologous recombination, we generated IRF7- KO A549 cells (Fig S5A) and reconstituted these cells with a stable transgene expressing C-terminally 3xFLAG-tagged cIRF7 or exIRF7 (Fig S5B). We confirmed these cell lines express similar levels of cIRF7 and exIRF7 (Fig S5B) and that both of these isoforms are retained in the cytoplasm in uninfected cells but transit efficiently to the nucleus upon pIC treatment (Fig S5C). Importantly, A549s expressing single isoforms of IRF7 fully recapitulate the exIRF7-specific IFNβ expression pattern observed in the Jurkat CRISPR clones (Fig S5D). In contrast to Jurkat cells, several IFNαs are induced by polyIC in A549 cells, and these are also specifically regulated by exIRF7(Fig S5D). Therefore, we conclude that exIRF7 uniquely induces transcriptional activation of type I IFN genes in response to PRR activation across diverse cell types and systems.

### exIRF7 preferentially dimerizes and forms specific complexes on the IFNβ promoter

To understand the molecular basis for the activity of exIRF7 on IFN-I expression and particularly IFNβ, we first asked if the promoter sequence for IFNβ was sufficient for the isoform-specific activation of transcription using luciferase reporter assays (Fig 3A). These experiments were done in HEK293 cells using the canonical stimuli Sendai virus (SENV) since SENV is a more robust stimuli of PRRs in HEK293 cells than pIC^18^. We found that introduction of the minimal core promoter of IFNβ (aka PRD31) displays modest induction by cIRF7 or exIRF7; however, transfection of the full IFNβ promoter reveals that exIRF7 exhibits more robust activation than cIRF7 in response to SENV (Fig 3A,B). These results suggest that the IFNβ promoter is sufficient for the differential activity of exIRF7 and cIRF7, and further confirm that the difference in activity between these proteins is observed across cell types and PRR stimuli.

**Figure 3:**
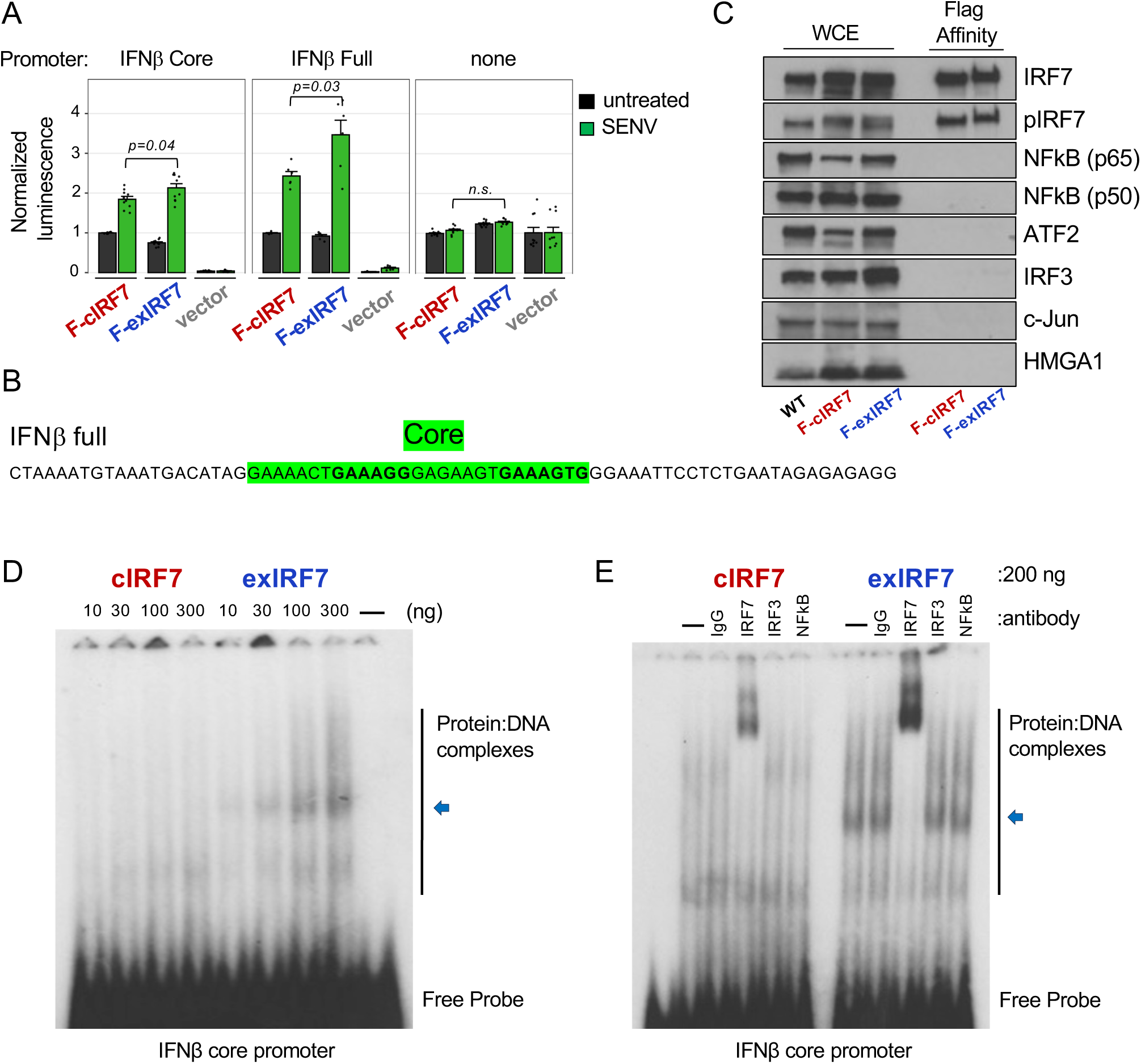
exIRF7 uniquely interacts with and preferentially enhances the activation of the IFNβ promoter. (**A**) Luciferase assay with cells co-expressing IRF7 isoforms, a NanoLuc internal control, and firefly luciferase under the control of an IFNβ promoter segment, after exposure to an innate immune trigger, sendai virus infection (500HAU/ml for 6hrs). Luciferase values are normalized to the NanoLuc control and shown relative to untreated cIRF7-expression. Significance shown was determined by unpaired t-test, each point is an independent biological replicate. (**B**) Sequence of IFNb promoter used in reporter assays and EMSAs (panels D-E). (**C**) Western blot analysis of Flag-affinity purified recombinant IRF7 proteins used in gel-shift assays. Proteins were purified from Jurkat cells stably expressing each isoform after treatment with pIC (10 µg/ml for 6 hours). The blot illustrates their relative expression levels, phosphorylation status, and absence of known IRF7 co-factors. (**D**) Representative native gel-shift assay (EMSA) using indicated amounts of IRF7 proteins from panel C incubated with a radiolabeled DNA oligo corresponding to core IFNβ promoter as shown in panel B (green sequence). Migration of free DNA probe and protein:DNA complexes is indicated. The blue arrows highlight the unique DNA:protein complex observed with exIRF7. Replicate gels with independent protein preps shown in Supplemental Fig S6. (**E**) EMSA as in panel D but with IRF7 protein held constant and antibodies to IRF7, known co-factors, and IgG control added.

Having shown that the IFNβ promoter is sufficient to reveal differential activity of exIRF7 versus cIRF7, we next investigated the binding of the IRF7 isoforms to the IFNβ promoter element by native gel analysis. We purified C-terminally flag tagged cIRF7 and exIRF7 from pIC treated Jurkat cells, and confirmed these were both phosphorylated and purified away from other established activators of IFNβ including IRF3 (Fig 3C). Consistent with previous studies^18,35^, both cIRF7 and exIRF7 bind efficiently to the core IFNβ promoter (Fig 3D). However, we observe a unique, more slowly migrating species with exIRF7 that is not observed with the cIRF7 isoform (Fig 3D, blue arrow; see also Fig S6). We confirmed that this unique species observed on the IFNβ promoter corresponds to IRF7 binding, and not some residual co-factor, as this species and others are only super-shifted by anti-IRF7 antibodies (Fig 3E).

Given that IRF7 alone forms distinctly migrating species on the native gels, and IRF7 is known to homodimerize, we asked if the larger EMSA species observed with exIRF7, might indicate a preference to form a dimer or higher order oligomer, even in the absence of DNA. Using Mass Photometry, we indeed observe that purified exIRF7 is primarily a dimer at concentrations used in these assays, while cIRF7 is predominantly monomeric (Fig 4A). Moreover, using a protein titration in the mass photometry, the Kd of dimerization for exIRF7 is calculated to be more than 6-fold tighter (i.e. higher affinity) than for cIRF7 (Fig. 4B).

**Figure 4:**
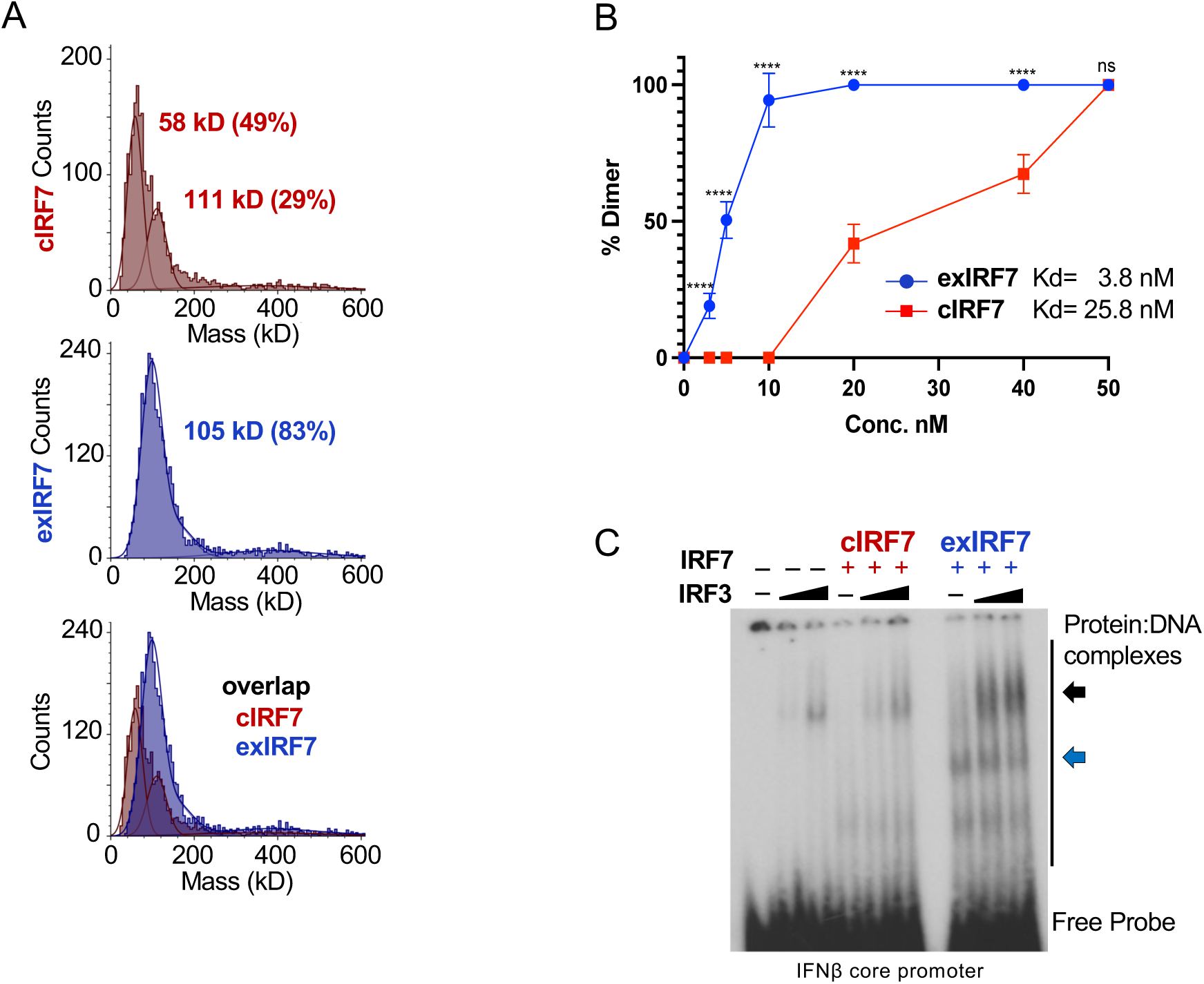
exIRF7 preferentially homodimerizes and binds cooperatively with IRF3. **(A)** Mass photometry traces of cIRF7 (top) and exIRF7 (middle). Mass of major populations are indicated as determined by fitting guassian curves as described in Methods. Bottom plot is the overlap of the individual traces**. (B)** Percent dimer of exIRF7 (blue) and cIRF7 (red) as calculated across a range of protein concentrations. Data points that differ significantly (p<0.001, unpaired student t-test) between the isoforms are indicated by ****. Kd calculated by curve fitting the data is shown. **(C)** Representative native gel-shift assay (EMSA) using 200 ng of purified IRF7 protein, and/or 20 or 40 ng of purified recombinant IRF3, incubated with a radiolabeled DNA oligo corresponding to core IFNβ promoter as shown in Figure 3. Migration of free DNA probe and protein:DNA complexes is indicated. The blue arrow highlights the unique exIRF7 complex, the black arrow indicates the IRF3 and IRF3/IRF7 complexes. A replicate gel with independent protein preps shown in Supplemental Fig S6.

In cells, IRF7 can function on the IFNβ promoter as both a homodimer as well as a heterodimer with IRF3^18,35^. Therefore, we also tested the ability of each of the isoforms of IRF7 to associate with IRF3 on the IFNβ promoter. Consistent with its increased tendency for dimerization, we also observe stronger cooperative binding between exIRF7 and IRF3 on IFNβ than cIRF7/IRF3 (Fig 4C, black arrow). Taken together, we conclude that the robust activity of exIRF7 in promoting expression of IFNβ reflects its enhanced ability to form the optimal enhancesome structure through cooperative binding with IRF3 and/or IRF7^35^.

### exIRF7 promotes inflammation and anti-viral immunity

Expression of IFNβ, and the resulting program of ISGs induced by IFN-I signaling, is central to the inflammatory response and anti-viral immunity. Too much interferon signaling can result in aberrant inflammation and autoimmunity, while too little interferon response increases susceptibility to infection. Therefore, the expression of IFNs must be tuned appropriately to maintain health. Given the unique ability of exIRF7 to drive strong IFNβ expression we hypothesized that cells may use alternative splicing of IRF7 to achieve appropriate IFNβ expression. A naturally occurring polymorphism in IRF7 in the human population (rs12290989 G/T) shows strong correlation with the splicing of IRF7 intron 1 by sQTL analysis (Fig 5A). Specifically, the major allele (GG) correlates with increased intron retention and expression of exIRF7, while the minor allele (TT) correlates with splicing of intron 1 and expression of cIRF7. Interestingly, presence of the minor allele is also associated with reduced expression of type I IFNs by plasmacytoid dendritic cells (pDC) in response to viral infection^36^, consistent with preferential expression of cIRF7 over exIRF7. Moreover, this same minor allele has been shown to be protective against the autoimmune disorder Lupus (SLE)^37^, as would be anticipated from a tendency to reduced IFNs and inflammation.

**Figure 5:**
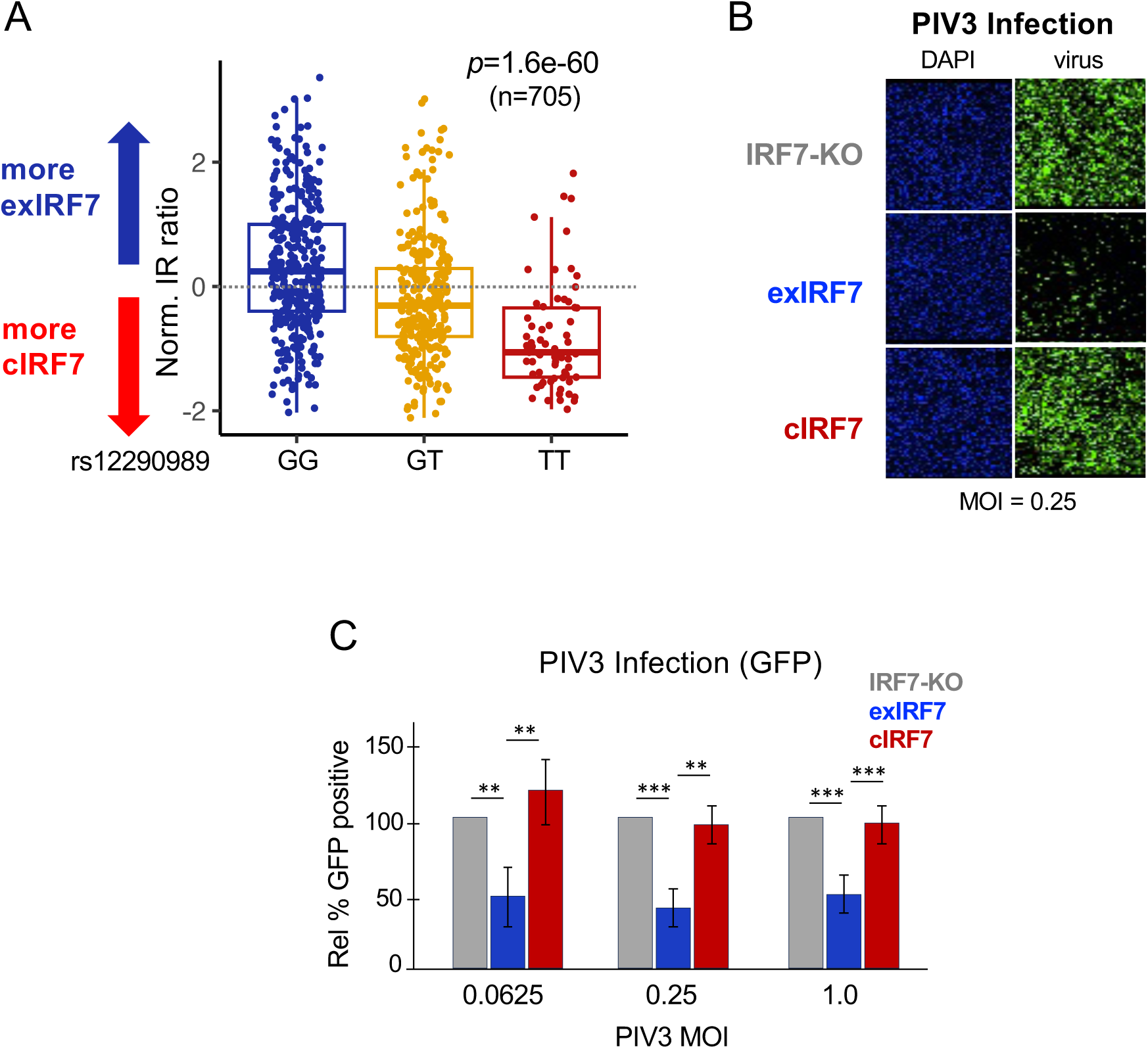
exIRF7 promotes interferon production and anti-viral immunity. (**A**) The CancerSplicingQTL database quantification of IRF7 intron 1 retention in different genotypes of SNP rs12290989. Each dot represents a single individual with the given genotype. Box plot represents average and distribution of intron retention. **(B)** Representative images from A549 cells with the indicated genotype infected for 24 hours with GFP-labeled PIV3 at MOI of 0.25. **(C)** Quantification of automated microscopy images as in panel B for PIV3 across multiple MOI of infection.

To more directly test the relevance of IRF7 isoform expression in the control of viral infections we used the A549 cells expressing single isoforms of IRF7 (Fig S4). Parainfluenza virus (PIV3) infects respiratory epithelial cells and has been shown to be highly sensitive to IFN-I signaling^38,39^. Strikingly, we observe that cells expressing exIRF7 are markedly more resistant to PIV3 infection than cells expressing no IRF7 or the cIRF7 isoform (Fig 5B), as would be expected from the increased expression of IFN-I in the exIRF7 cells (Fig S4). We further substantiated the difference in PIV3 viral infection in exIRF7 cells relative to cIRF7 across multiple MOIs by quantitative microscopy (Fig 5C). These data confirm the physiologic relevance of the differential activity of the IRF7 isoforms in initiating an anti-viral immune response.

## Discussion

The role of the transcription factors IRF7 and IRF3 in activating the expression of interferons following innate immune triggers has been studied for several decades, yet the intricacies regarding how this process is regulated remain to be fully uncovered. Here we demonstrate a previously unrecognized mechanism by which alternative splicing of IRF7 tunes the transcriptional activation of type I IFNs and subsequent downstream ISGs. Specifically, we show that the N-terminally extended form of IRF7 (exIRF7), which is encoded by retention of the first intron within the coding region of the human IRF7 gene, has greater activity on the IFNβ promoter due to increased propensity to dimerize. The increased activity of exIRF7 on the IFNβ promoter, in turn, induces higher levels of IFN and ISGs to repress viral infection compared to the canonical (cIRF7) isoform.

Most previous work on IRF7 has focused on the cIRF7 isoform, due in part to the fact that cIRF7 is the only isoform expressed in mice, as the murine IRF7 gene lacks the in-frame AUG which initiates translation of the exIRF7 isoform. Indeed, the potential to encode exIRF7 appears to be restricted to Old world monkeys and Apes (which includes humans), suggesting evolutionary divergence ∼21.9 million years ago. While mice do not encode the exIRF7 isoform, alternative splicing does regulate murine IRF7 activity through intron 4 retention to attenuate translation^31^. We do not observe this intron 4 mechanism in humans, at least in the conditions and cells surveyed in this study. Furthermore, the splicing factor SRSF7 enhances murine IRF7 transcription through recruiting chromatin remodeling enzymes^40^. Therefore, it is possible that tuning of the interferon response through the activity of the splicing machinery has been conserved throughout evolution, with different species adopting distinct mechanisms to establish this connection.

While the existence of the exIRF7 isoform, and its ability to support transcription of IFNα^33^ was previously established, comparisons between the activity of exIRF7 and cIRF7 have not been explored. Here we directly compare the activity of these isoforms in three different cell lines, studying the endogenous genes and reporter assays. In all cases, exIRF7 exhibits increased transcriptional activation of a subset of known target promoters, including, critically, those driving type I IFN genes. The discovery of enhanced activity of exIRF7 provides critical new insight into the tuning of the interferon response. Nevertheless, our observations are consistent with previous work in that upon ectopic expression (Fig 3A), or after IFN stimulation (Fig 2E), the high levels of cIRF7 indeed compensate for the reduced transcriptional activity and activate the interferon pathway.

The formation of higher order complexes on the IFNβ promoter is a critical step in nucleating the enhanceosome complex to drive transcription^35^. Mechanistically, we find that the enhanced ability of exIRF7 to activate transcription correlates with its increased ability to dimerize compared to cIRF7. The increased abundance of higher order exIRF7 forms, both on its own and in complex with IRF3, as compared to cIRF7, is consistent with the increased IFNβ and ISG expression in cells expressing exIRF7. Previous studies demonstrated that dimerization of cIRF7 with IRF3 is mediated by the C-terminus of both proteins, which is identical in both IRF7 isoforms. We consider it likely that the extended N-terminus of exIRF7 makes contact with DNA, thus increasing the formation of DNA-bound complexes, as the N-terminus is immediate adjacent to the DNA binding domain. Alphafold does not predict any stable structure of the N-terminal peptides in either cIRF7 or exIRF7 and the N-terminus of cIRF7 was truncated in the published structure studies of IRF7 bound to DNA^35^. Future studies, requiring extensive structural analysis, will thus be required to provide insight into how the N-terminal extension of IRF7 enhances dimerization and function.

Finally, we demonstrate that exIRF7 not only increases transcriptional activation of the interferon response, but also enhances antiviral defense as demonstrated by reduced susceptibility to viral infection. Of note, many human immune cell types that control viral infections including monocytes, memory B and T cells, and Tregs preferentially express exIRF7 (Fig 1B), suggesting that these cells use exIRF7 expression to promote initial responsiveness of the immune system. Conversely, increased expression of cIRF7 following immune activation of monocytes may promote downregulation of the interferon response to prevent hyperinflammation. We also found that viral infection itself can alter splicing of IRF7, reducing the relative expression of exIRF7, which may be one of the many mechanisms viruses use to counter the interferon response and enhance their replication. Altogether, we propose that alternative splicing of IRF7 serves as a regulatory mechanism to toggle between initial activation and subsequent homeostasis of the innate immune system.

## Acknowledgements

We wish to thank the Penn Human Immunology Core (RRID SCR_022380) for providing primary monocytes, the CDB Microscopy Core (RRID SCR_022373) for microscope time, and Dr. Kushol Gupta and the Johnson Foundation Biophysics and Structural Biology Core (RRID SCR_022414) for instruction and use of the Mass Photometer. We also thank Mark Dittmar for RNA from viral infected cells, and Drs. Simon Boudreault (Penn) and Yunsum Nam (UT Southwestern) for advice and suggestions on this work. This work was funded by R35 GM118048 to KWL and R01 AI150246, R01 AI152362 and R01 AI140539 to SC.

## Author Contributions

K.W.L. conceived of and directed the project with assistance from S.C., A.P. and M.F. A.P. carried out all the Jurkat cell work and biochemistry, M.F. carried out all the RNA-Seq analysis, O.H. generated and analyzed the A549 data, S.M.P. carried out the viral work with PI3V, R.M. assisted with RT-PCR analysis, M.Q.V. did the GTEX analysis of IRF7 splicing across tissues prior to publication of the MAJIQlopedia method, Q.L. did the sQTL analysis. K.W.L., S.C., A.P. and M.F. wrote and edited the manuscript with comments from all the authors.

## Methods

### Experimental Model Details

Wild-type, CRISPR-edited, and cDNA-overexpressing stable Jurkat (JSL1) cells were cultured in RPMI medium (Corning, 10-040-CV) supplemented with 5% heat-inactivated fetal bovine serum (FBS) (Gibco, A56697-01), and 2 mM penicillin-streptomycin (Mediatech, MT-30-002-Cl). The cells were maintained at 37°C in a humidified incubator with 5% CO₂. A549 and HEK293T cell lines were grown in DMEM (Corning, 10-013-CV) supplemented with 10% FBS, under the same conditions as those described for JSL1 cells. Human primary monocytes were obtained from the Human Immunology Core at the University of Pennsylvania and were maintained in RPMI with 10% FBS. To stimulate innate immune responses, the cells were adjusted to a density of 1 × 10⁶ cells/ml and incubated either in medium alone or with the designated treatments for the required duration.

### RT-PCR

RNA was extracted from Jurkat cells, primary monocytes, or A549 cells using TRIzol (Invitrogen, 15596018) in accordance with the manufacturer’s instructions. Low-cycle radioactive RT-PCR was conducted and analyzed as previously detailed (Lynch and Weiss, 2000; Ip et al., 2007; Rothrock et al., 2003). RNA was reverse transcribed using MMLV reverse transcriptase (Thermo, 28025013). The primer sequences for IRF7 splicing analysis are listed in the Key Resource Table. Three independent RT-PCR reactions were performed with 500 ng of RNA per reaction, and samples were resolved on 5% denaturing polyacrylamide gels (PAGE) using formamide buffer. Radiolabeled 32P cDNA products were detected via densitometry with a Typhoon PhosphorImager (Amersham Biosciences) and quantified using ImageQuant software.

### Western Blots

Whole cell lysate was collected with RIPA lysis buffer (150 mM NaCl, 0.5% Sodium Deoxycholate, 1% NP-40, 0.1%SDS, 25mM Tris (pH 7.4)) and proteins were quantified by Bradford assay. 20 µg of total protein lysates were loaded into 10% 37.5:1 bis-acrylamide SDS-PAGE gels. SDS-PAGE gels were transferred onto PDVF membranes and targeted proteins were visualized with a chemiluminescence system and subsequent imaging with an X-ray developer. Antibodies used to detect protein expression levels are as follows: IRF7 (Abcam, ab238137), pIRF7 (Cell Signaling, 5184S), FLAG (Cell Signaling, 14793S), IRF3 (Cell Signaling, 11904S), NFkB-p65 (Cell Signaling, 8242T), NFkB-p50 (Cell Signaling, 12540S), ATF2 (Cell Signaling, 35031S), c-Jun (Cell Signaling, 9165T), HMGA1 (Cell Signaling, 12094S), and GAPDH (Cell Signaling, 2118S), anti-rabbit HRP-linked secondary antibody (Cell signaling, 7074S). All the primary antibodies were used at a 1:1000 dilution and the secondary antibody was used at a 1:3000 dilution.

### CRISPR designs and cloning

To generate individual cell clones that express either isoform of IRF7, we used CRISPR/Cas9- based genome editing in Jurkat cells to alter the genomic elements that encode for the regulated alternative splicing events. Two custom sgRNAs (sgRNA#1 and sgRNA#2) were created to target intronic regions or the 5’UTR that surround genomic elements that encode for the alternative splicing event of interest. To knockout IRF7, two sgRNAs (sgRNA#2 & sgRNA#3) were designed targeting the introns adjacent to the constitutive exon of interest. The sgRNAs designed sequences are listed in the Key Resources Table. The sgRNAs were designed using the Broad Institute CRISPick tool (https://portals.broadinstitute.org/gppx/crispick/public) and cloned into the pSpCas9(BB)-2A-GFP (PX458) plasmid (Addgene, 48138) containing the S. pyogenes Cas9 (SpCas9), GFP and U6 promoter sequences. For cIRF7, a repair template was constructed to fuse exons 1 and 2, mimicking the splicing outcome when Intron 1 is skipped. To generate exIRF7, the repair template included a mutation in the 5’ splice site (5’SS), changing the splice donor “G” of the 5’SS to “C” (AGG to CGC, R-R), thereby blocking splicing and resulting in Intron 1 retention. Both repair templates were synthesized as gBlocks by IDT. Upstream and downstream homology arms (∼600 bp each) were PCR-amplified (with KAPA PCR kit, Roche, KK2602) and assembled into the pUC19 vector backbone using the NEBuilder HiFi DNA Assembly Kit (NEB, E2621L). The sequences for the repair templates, primers for homology arm amplification, and gBlocks are listed in the Key Resource Table.

For exogenous IRF7 expression, the cIRF7 (NM_001572; 504 amino acids; GenScript Clone ID: OHu24955) and exIRF7 (NM_004031; 517 amino acids; GenScript Clone ID: OHu25038) human ORF cDNAs were cloned into the pEFneo vector. A 3xFLAG tag (DYKDHDGDYKDHDIDYKDDDDK) was added to the C-terminus of each construct for downstream applications. The final engineered plasmids contained the 3×FLAG tag fused to the C-terminal ends of the IRF7 isoforms.

### RNA-Seq

CRISPR-edited clones were treated with either high molecular weight (HMW) polyinosinic-polycytidylic acid (polyIC) (Invivogen, tlrl-pic) at a final concentration of 1μg/ml or recombinant interferon-alpha 2a (IFN-Alpha 2a) (PBL Assay Science, 11100-1) at a final concentration of 500 U/ml for 8 hours. Following an 8-hour treatment with either polyIC or IFN-Alpha 2a, cells were collected, and RNA was isolated using TRIzol reagent according to the manufacturer’s instructions. RNA integrity number (RIN) was measured with the Agilent bioanalyzer, and all samples had a RIN >8.0. RNA-sequencing libraries were generated by and sequenced by GeneWiz (Azenta life sciences) at a depth of 20 million reads per sample. The libraries were poly(A) selected (nonstranded) and paired-end sequenced at a 150 bp read length. Raw fastq files were trimmed with bbduk 38.79 (www.osti.gov/biblio/1241166) and then counted with salmon 1.10.0^41^ in quant mode with the settings “--validateMappings --rangeFactorizationBins 4 –seqBias --gcBias --recoverOrphans”. Reads were aligned to Ensembl Homo sapiens hg38 primary cDNA transcriptome (annotation version 110). Differential expression analysis was performed in R using Tidyverse packages for data management and processing (doi:10.21105/joss.01686). Transcript counts from salmon were collapsed to the gene level using tximport^42^ and differential abundance was determined using DESeq2^43^. Differential expression analysis was quantified by DESeq2. In Figure 2, differential expression of a gene is considered significant with a pValue < 0.05, log2 fold-change > 1 (stimulated/unstimulated) and base mean reads > 50. The RNA-sequencing data generated for this study is available in GEO (<u>GSE285352</u>).

### ISRE analysis

Transcriptional start sites (TSS) were extracted from Ensembl using the biomaRt package^44^. Next, the regions around the TSS were cross referenced against the JASPAR database^45^ to note the location of all ISRE (i.e., predicted STAT1::STAT2) binding sites. To generate the logos, the ISRE sites from each gene group were extracted and visualized using ggseqlogo^46^.

### RT-qPCR

Total RNA was extracted from cells using TRIzol, following the manufacturer’s protocol (Invitrogen, 15596018). cDNA was synthesized from 250 ng of RNA using oligo(dT) primers (Invitrogen, 184118012) and MMLV reverse transcriptase (Thermo, 28025013). Quantitative PCR was performed using SYBR qPCR Master Mix (Applied Biosystems, A25742) on the QuantStudio 6 Flex Real-Time PCR System (Applied Biosystems). Data acquisition was managed with QuantStudio 12K Flex software version 1.3 (Applied Biosystems) and analyzed using the ΔCT method. Actb, encoding β-actin, was used as the reference gene for normalization. The sequences of the gene-specific primers utilized for PCR are listed in the Key Resource Table.

### Construction of plasmids and transfection

All transfections in JSL1 cells were done by electroporating 10–20 million cells. To generate cIRF7 and exIRF7 cells, Jurkat cells were co-transfected with PX458-sgRNA plasmids targeting intronic regions flanking the splicing event and pUC19-repair template plasmids containing homologous arms and repair templates (5 µg each). To generate IRF7 knockout (KO) cells, PX458-sgRNA plasmids targeting the introns flanking the constitutive exon of interest were co-transfected into cells (5 µg each). After electroporation, edited Jurkat cells were rested for 48hrs and sorted into 96-well plates (FACS Jazz) by gating on doublet exclusion and GFP+ expression. One to three cells were deposited per well to allow the growth of single-cell colonies. Individual clones were screened by PCR amplification of the targeted genomic regions, analyzed for band size on a 1.5% EtBr agarose gel, confirmed through genomic DNA sequencing, and validated by Western blot analysis. All Jurkat cells were maintained in RPMI medium supplemented with 5% heat inactivated FBS. Stable expression of cIRF7 and exIRF7 with C-terminal 3xFLAG tags was achieved by integrating pEFneo cDNA expression constructs into JSL1 cells. Linearized plasmids (10µg) were electroporated into 10–20 million cells, and neomycin was used as a selective marker for stable integration.

### Electrophoretic Mobility Shift Assay

Jurkat cells stably expressing individual IRF7 isoforms tagged with FLAG were stimulated with 1 µg/mL poly(I:C) for 6 hours prior to harvesting. IRF7 protein was purified using anti-FLAG M2 affinity agarose gel beads (Sigma, A2220) via FLAG-pulldown. The FLAG-purified recombinant IRF7 proteins were analyzed for DNA binding via gel-shift assays (EMSA) using a 32P-labeled double-stranded oligonucleotide corresponding to the PRD3,1 region of the IFNβ promoter (5’- GAAAACTGAAAGGGAGAAGTGAAAGTG-3’). The EMSA binding buffer consisted of 20 mM Tris-HCl (pH 7.5), 1 mM EDTA, 20 mM KCl, 1 mM MgCl₂, 10% glycerol, 5 mM DTT, 0.5% NP-40, 10 µg of BSA, and 62.5 µg/mL Poly(dI-dC) (Thermo Scientific, 20148E) to minimize nonspecific binding. For each binding reaction, 40,000 cpm of the radiolabeled probe was added. Reactions were incubated at 30°C for 20 minutes, then immediately placed on ice. For supershift assays, recombinant proteins in the binding buffer were pre-incubated with 0.2 µg of antibodies against IRF7 (Abcam, ab238137), IRF3 (Cell Signaling, 11904S), NF-kB (Cell Signaling, 8242T), or rabbit IgG secondary antibody (Cell Signaling, 7074S) prior to adding the radioactive probe. Following the binding reactions, protein-DNA complexes were separated on 5% acrylamide/bis-acrylamide gels (29:1 cross-link ratio, Bio-Rad, 1610156) prepared in 0.5× TBE buffer. Gels were run at 200 volts for 2 hours, dried, and exposed to film at -80°C for 2 hours. The complexes were visualized using an X-ray developer.

### Mass Photometry

For Mass Photometry experiments, we used the same FLAG-purified batch of cIRF7 and exIRF7 proteins used in the previously described gel-shift assays. Measurements were carried out in silicone gaskets placed on mass photometry coverslips (MP consumables, Refeyn RD501078). Immediately prior to mass photometry measurements, protein stocks were added to a buffer (20 mM Tris-HCl (pH 7.5), 20 mM KCl, 1 mM MgCl₂, and 5 mM DTT). The reactions were incubated at 30°C for 20 minutes, then immediately placed on ice. The final working concentrations of protein was 20 nM. For each measurement, 20µL of sample (protein+ buffer) was added to a silicone gasket placed on a coverslip. Following autofocus stabilization, movies of either 60 or 120s duration were recorded. Data acquisition was started ≤5 s after the addition of proteins. Each sample was measured at least three times independently (*n* ≥ 3). All experiments were conducted using the same mass photometry instrument, the TwoMP mass photometer (Refeyn Ltd, Oxford, UK). Data acquisition was performed using AcquireMP software, and subsequent analysis was carried out with DiscoverMP (Refeyn Ltd).

### Immunofluorescence

A549 cells expressing IRF7 isoforms were fixed with 4% formaldehyde for 20 minutes at room temperature and then washed with PBS to remove residual formaldehyde. Permeabilization was performed in a permeabilization buffer (PBS containing 0.15% Triton-X100) for 5 minutes at room temperature and then washed twice with PBS. Blocking was performed using a blocking buffer (PBS containing 10% normal goat serum (NGS)) for 20 minutes at room temperature. Primary antibodies were diluted in blocking buffer and incubated with the cells overnight at 4°C. Following primary antibody incubation, cells were washed three times with PBS. Fluorophore-conjugated, species-specific secondary antibodies were diluted in blocking buffer and applied to the cells for 1 hour at room temperature. Cells were then washed three times with PBS and stained with Hoechst 3342 in PBS for fifteen minutes at room temperature. Cells were then washed three additional times with PBS and visualized using the Zeiss LSM 980 Confocal Microscope. Image analysis was performed with the FUJI imaging platform.

### Luciferase Reporter Assay

HEK293T cells were seeded at a density of 1 × 10^4^ cells/well in 96-well plates containing DMEM supplemented with 10% FBS. Cells were co-transfected for 48 hours with 100ng of plasmids encoding 40ng of IRF7 isoforms (pEFneo-cIRF7 or pEFneo-exIRF7) or an empty vector, 10ng of pNL1.1.TK [Nluc/TK] to drive NanoLuc activity, and 50ng Firefly luciferase constructs with IFN-β promoter segments cloned upstream of the minimal promoter in the pGL4.26 luc2-minP- Hygro vector, or PsiCHECK2 (as a control). Transfections were performed using Lipofectamine 2000 (Thermo Fisher, 11668027) according to the manufacturer’s protocol. For assays involving Sendai Virus (SeV) stimulation, cells were infected with 500 HA units/mL of SeV (avsbio, 10100774) in the media for 8 hours. Luciferase activity was measured on a Tecan Spark multimode plate reader using the Nano-Glo® Luciferase Assay System (Promega, N1110) as per the manufacturer’s instructions. Luciferase activity was normalized to uninfected cells expressing pEFneo-cIRF7.

### Viral infections and quantitation

For automated microscopy of viral infections, human parainfluenza virus type 3 (HPIV3-GFP) was propagated in LLC-MK2 cells. A549 cells (IRF7-KO, exIRF7 and cIRF7) at a density of 2×10^4^ per well were plated in 96-well black tissue culture plate and infected with HPIV3-GFP at different MOIs for 24 hour. Cells were fixed using 4% formaldehyde for 15mins at room temperature then washed with PBS three times and blocked in blocking buffer (PBST (0.1% TritonX-100) + 2% Bovine Serum Albumin (BSA)) and stained with Hoechst 3342 for nuclei in blocking buffer for 1h. Cells were washed three times with PBS-T and imaged on a Molecular Devices ImageXpress Micro 4 imaging system. Images were quantified using Molecular Devices MetaXpress (version 6) modules and percentage of infected cells was calculated.

For viral RNA, A549 cells (IRF7-KO, exIRF7 and cIRF7) at a density of 2×10^5^ were plated in 12- well format and infected with HPIV3-GFP for 4h at an indicated MOI. After 4hpi, viral inoculum was removed, washed with PBS and cells were replenished with fresh media. Cells were lysed using Trizol reagent and viral supernatants were collected following 24 hour post infection. Viral RNA was assessed via RT-qPCR using HPIV3 primers as indicated in the Key Resource Table. The relative expression of viral RNA were calculated using standard curve method and normalized to 18s ribosomal RNA.

