## Supplemental Figures and Legends for "Alternate isoforms of IRF7 Differentially Regulate Interferon Expression to Tune Response to Viral Infection"

### Figure S1: Panthi et al

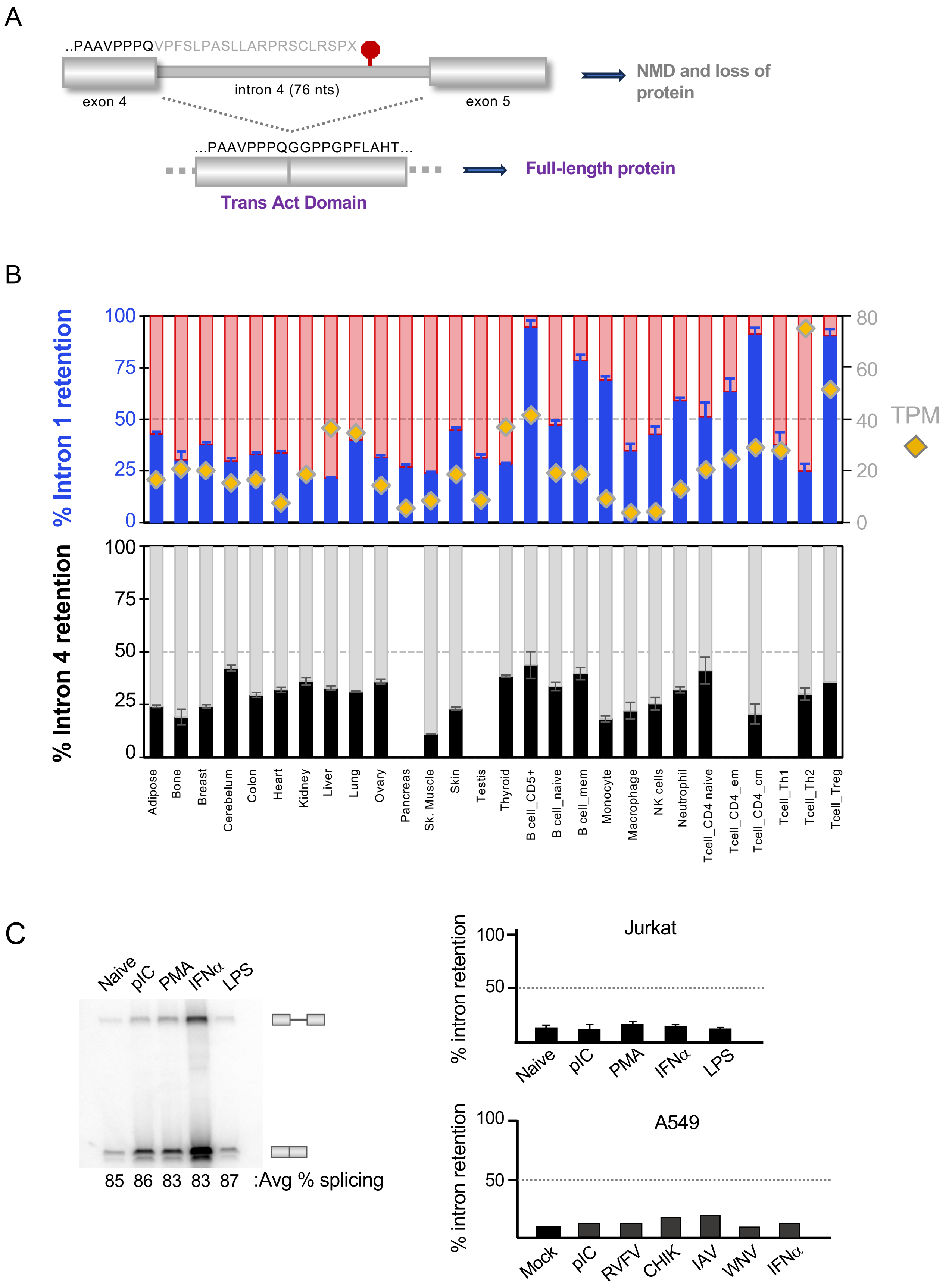

**<u>Figure S1</u>: Innate immune activation modulates IRF7 intron 4 retention in mice, but not in humans, unlike intron 1.**

**(A)** Schematic depicting IRF7 intron 4 retention, which leads to a premature stop codon in mice. (**B**) GTEx data illustrating the percent of total IRF7 mRNA that includes intron 4 across various cell types (black bars in the graph represent the intron 4-retained isoform), plotted in line with the retention of IRF7 intron 1 in the same cell types as from Figure 1B. Gold diamonds indicate that total abundance (TPM) of IRF7 in the indicated cell types. (**C**) RT-PCR analysis of IRF7 intron 4 alternative splicing in Jurkat and A549 cells in response to different immune triggers. (left) raw data from a subset of samples. (right) quantification of all conditions. *Related to Figure 1*.

### Figure S2: Panthi et al

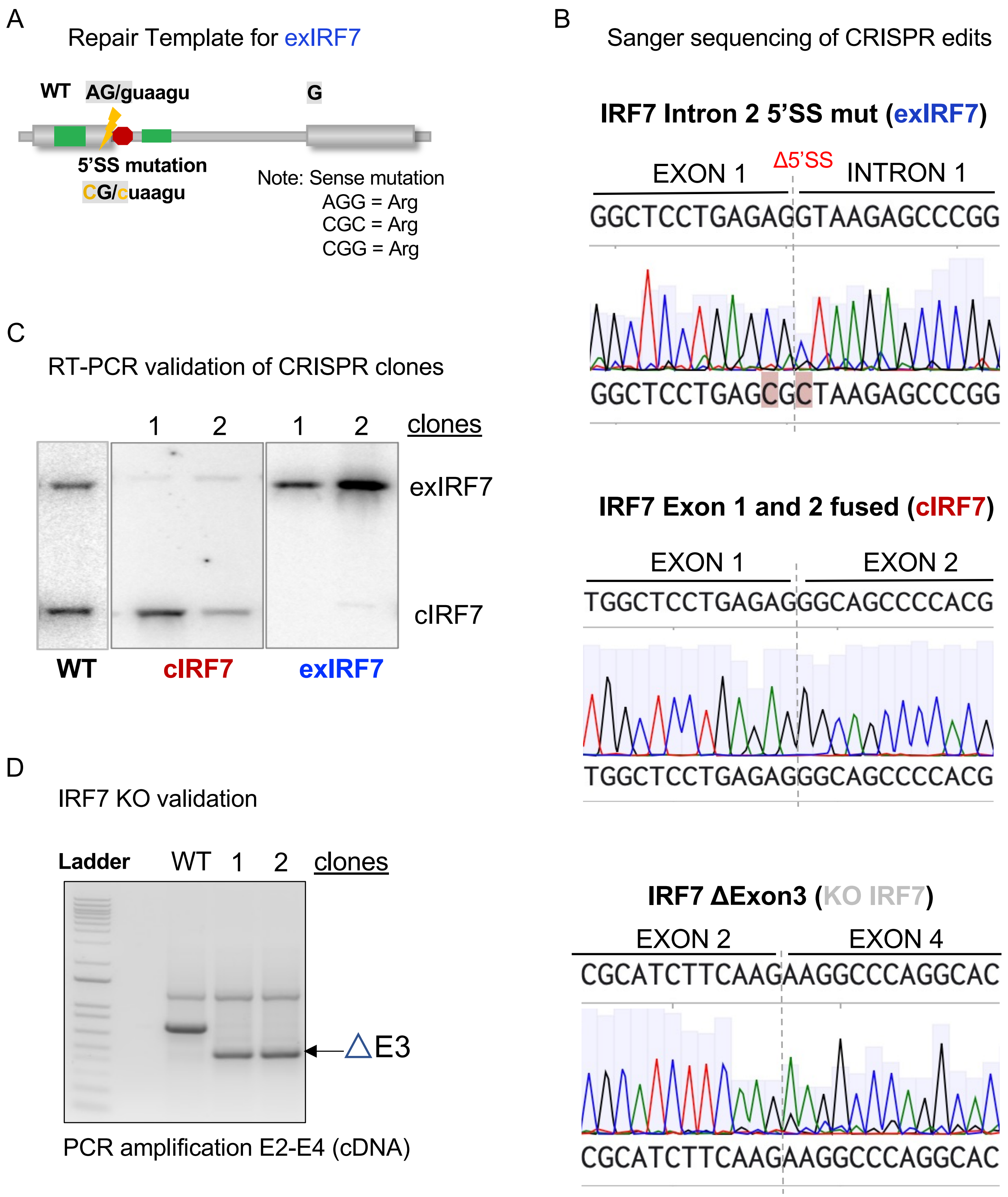

**Figure S2: Design and validation of CRISPR-based genome editing.**

(**A**) Schematic representation of the CRISPR repair template designed to generate exIRF7 with a 5' splice site mutation. The nucleotide alterations leading to the sense mutation are highlighted. (**B**) Sanger sequencing of genomic DNA extracted from the corresponding CRISPR-engineered cell lines. The sequencing peaks indicate the edited or deleted regions within the IRF7 genome. (**C**) Validation of CRISPR clones for each IRF7 isoform at the RNA level using RT-PCR. (**D**) Confirmation of IRF7 knockout clones at the RNA level through cDNA-PCR amplification. *Related to Figure 2.*

### Figure S3: Panthi et al

A

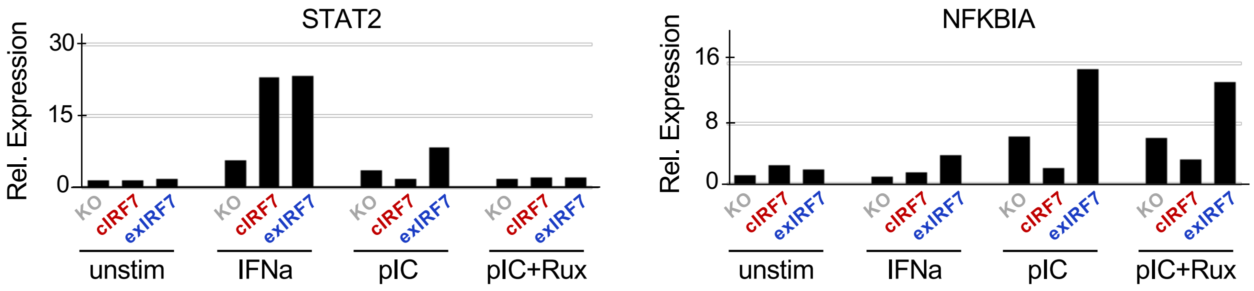


B

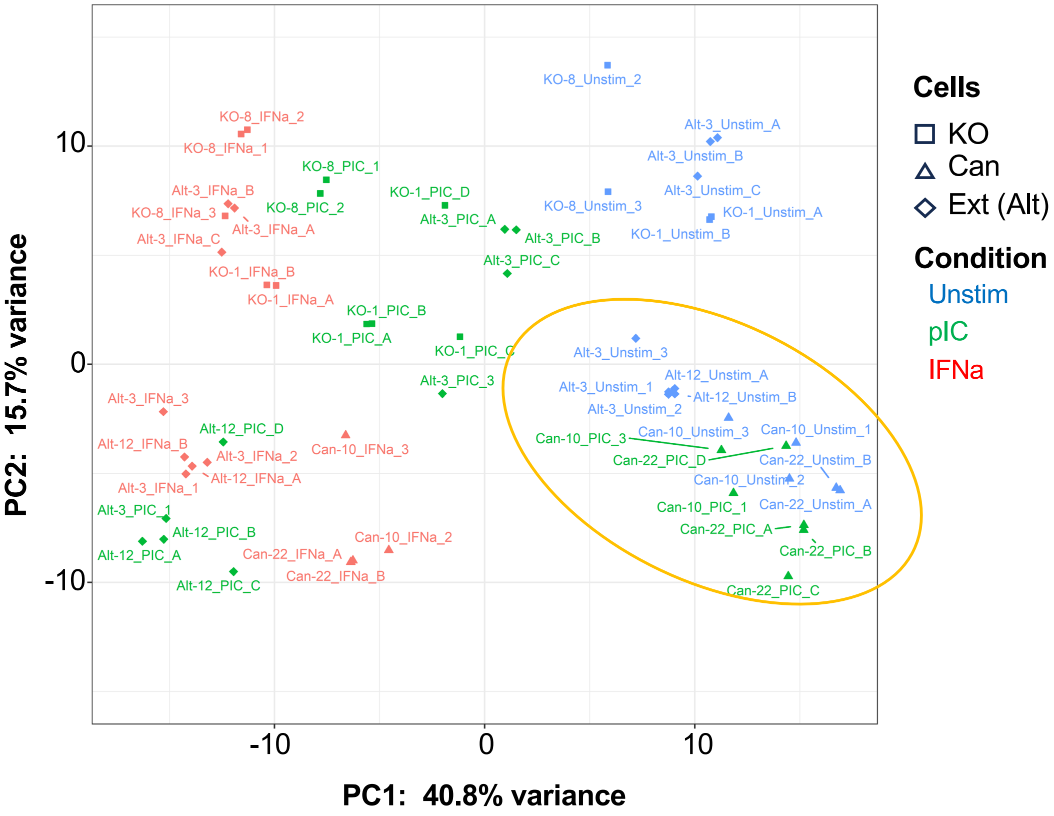

**<u>Figure S3</u>: PCA plot and additional validation of RNA-seq data**

(**A**) RT-qPCR analysis of additional differentially expressed genes (DEGs) from RNA-seq (n=3, mean ± SD) under conditions of IFNα and pIC, with and without ruxolitinib treatment. (**B**) Principal component analysis (PCA) plot of replicates illustrates gene expression in IRF7 knockout (KO), cIRF7, and exIRF7 cells following induction with pIC and IFNα. Different colors represent various stimulation conditions. The yellow circle in the bottom right highlights the clustering of unstimulated and pIC-treated cIRF7 cells alongside unstimulated exIRF7 and IRF7 KO cells. *Related to Figure 2*.

### Figure S4: Panthi et al

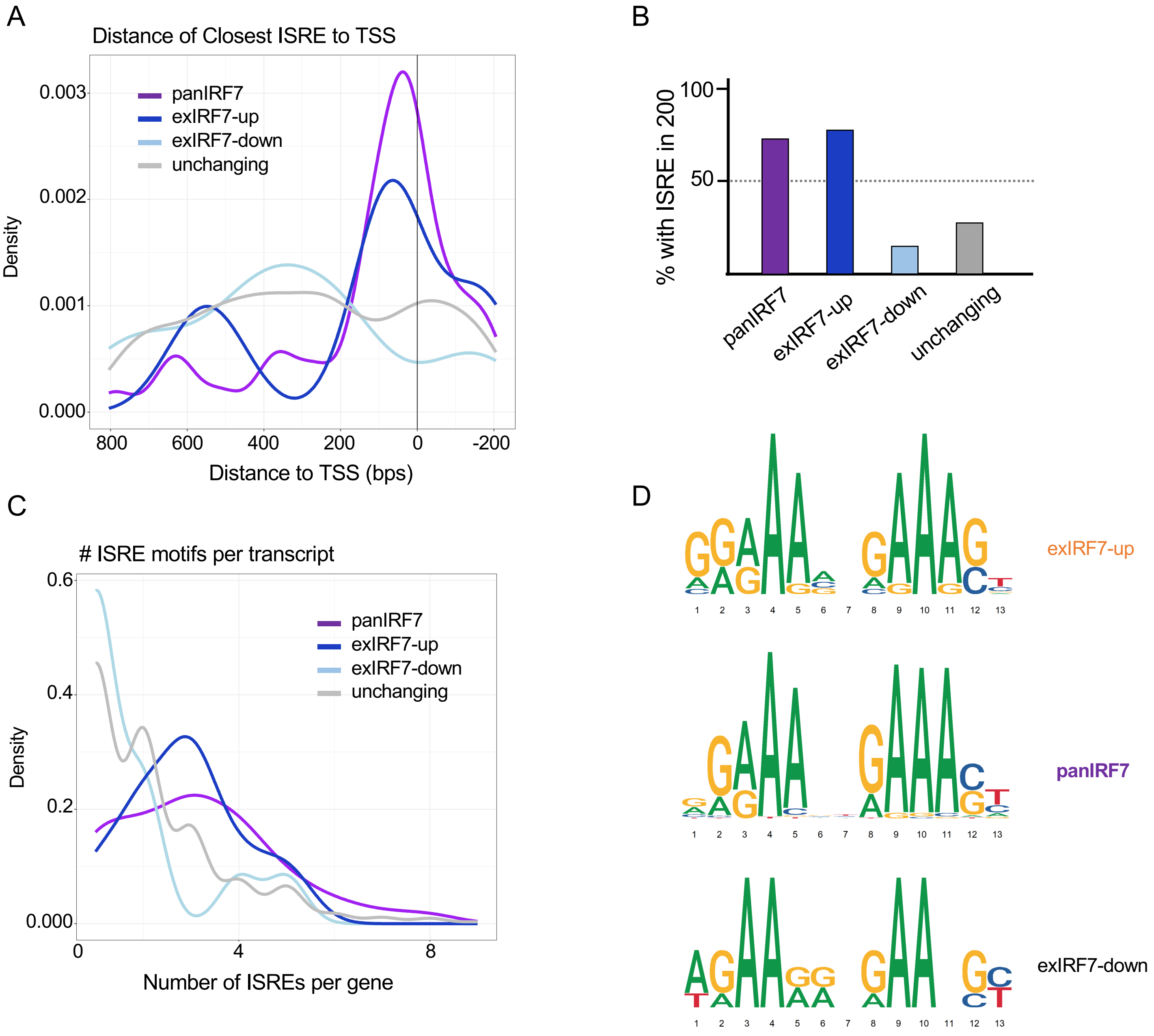

**<u>Figure S4</u>: Analysis of promoter sequences of genes expressed in distinct patterns in RNA-Seq data from Figure 2.**

(**A**) Density of IRF consensus sequences (ISRE) relative to transcription start site (TSS) across genes defined in Figure 2 to be enhanced by either IRF7 isoform in response to IFN treatment (panIRF7), uniquely enhanced (dark blue) or repressed (light blue) in exIRF7 cells in response to pIC, or unresponsive controls. **(B)** Percentage of genes in each category from panel A that have an IRF consensus sequence (ISRE) within 200 nts of the TSS. **(C)** Number of ISREs upstream of TSS in genes defined in panel A. **(D)** Most ISRE-like consensus sequence in 200 nts upstream of TSS in each gene category from panel A. *Related to Figure 2*.

### Figure S5: Panthi et al

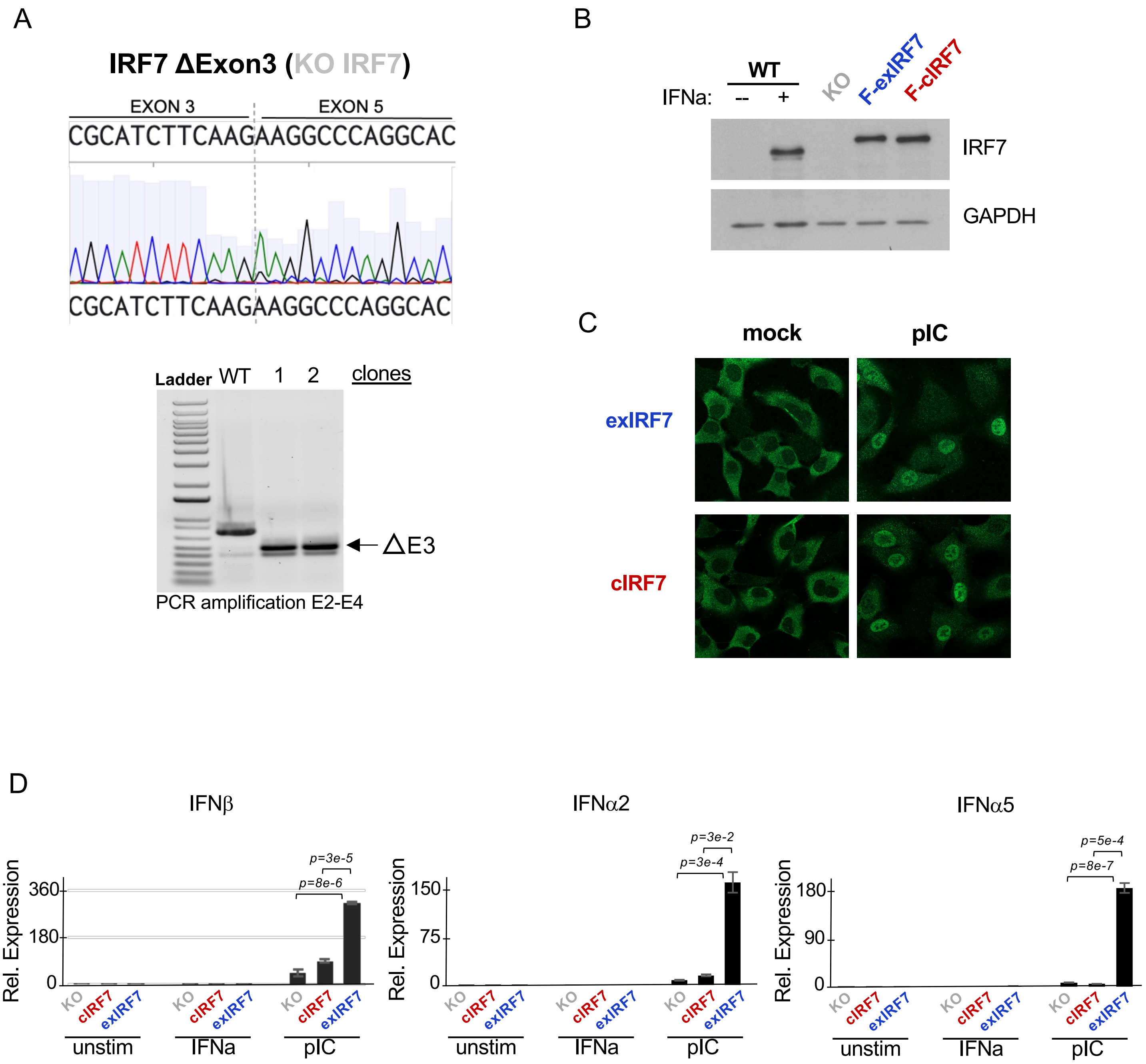

**<u>Figure S5</u>: Generation and characterization of A549 cells that express single isoforms of IRF7.**

(**A**) Sanger sequencing of genomic DNA extracted from IRF7 KO A549 cell lines (top). Confirmation of IRF7 knockout clones at the DNA level by PCR amplification (bottom). **(B)** Western blot of stable expression of C-terminal 3xFlag-tagged IRF7 isoforms (F-IRF7) in A549 clones relative to endogenous expression in IFN-treated cells. Top blot probed with antibody to IRF7, bottom blot probe with antibody to GAPDH as loading control. **(C)** Immunofluoresence with antibody to Flag-tag showing localization of tagged IRF7 isoforms before and after pIC treatment of A549 clones from panel B. **(D)** qPCR analysis of type I IFN gene expression in clones from panel B. *Related to Figure 2*.

### Figure S6: Panthi et al

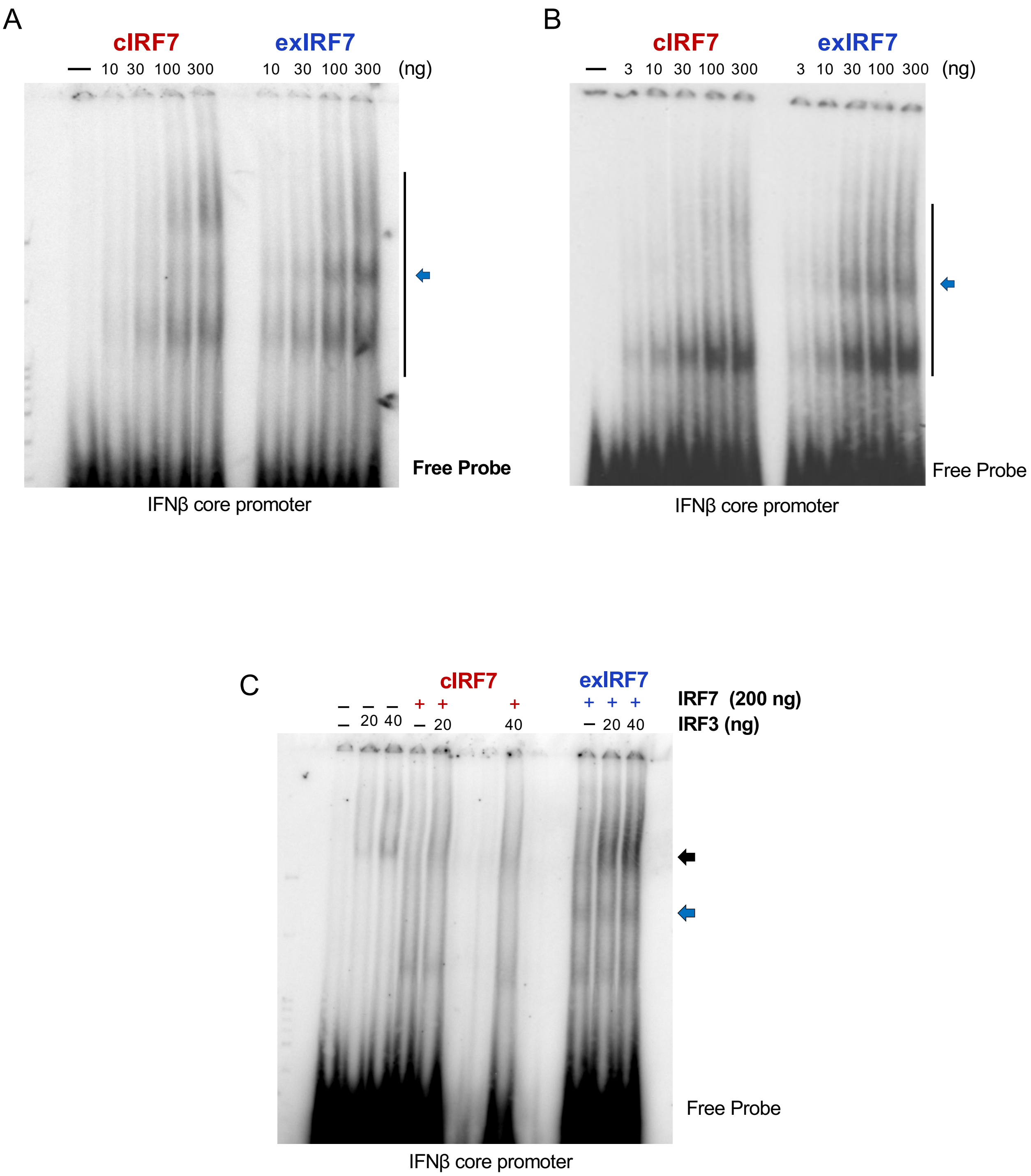

**Figure S6: Replicate EMSA of IRF7 protein isoforms with IFNβ core promoter.**

**(A-B)** Native EMSAs as in Figure 3D but done with independent preparations of purified IRF7 proteins, demonstrating reproducibility of unique exIRF7 species. **(C)** Native EMSA as in Figure 4C but with independent preparations of purified IRF7 proteins. *Related to Figures 3 and 4*.
